# Evaluation of methods to assign cell type labels to cell clusters from single-cell RNAsequencing data

**DOI:** 10.1101/562082

**Authors:** J. Javier Díaz-Mejía, Elaine C. Meng, Alexander R. Pico, Sonya A. MacParland, Troy Ketela, Trevor J. Pugh, Gary D. Bader, John H. Morris

## Abstract

Identification of cell type subpopulations from complex cell mixtures using single-cell RNA-sequencing (scRNA-seq) data includes automated computational steps like data normalization, dimensionality reduction and cell clustering. However, assigning cell type labels to cell clusters is still conducted manually by most researchers, resulting in limited documentation, low reproducibility and uncontrolled vocabularies. Two bottlenecks to automating this task are the scarcity of reference cell type gene expression signatures and that some dedicated methods are available only as web servers with limited cell type gene expression signatures. In this study, we benchmarked four methods (CIBERSORT, GSEA, GSVA, and ORA) for the task of assigning cell type labels to cell clusters from scRNA-seq data. We used scRNA-seq datasets from liver, peripheral blood mononuclear cells and retinal neurons for which reference cell type gene expression signatures were available. Our results show that, in general, all four methods show a high performance in the task as evaluated by Receiver Operating Characteristic curve analysis (average AUC = 0.94, sd = 0.036), whereas Precision-Recall curve analyses show a wide variation depending on the method and dataset (average AUC = 0.53, sd = 0.24). CIBERSORT and GSVA were the top two performers. Additionally, GSVA was the fastest of the four methods and was more robust in cell type gene expression signature subsampling simulations. We provide an extensible framework to evaluate other methods and datasets at https://github.com/jdime/scRNAseq_cell_cluster_labeling.

## Introduction

During the last five years a number of single-cell sequencing technologies have been developed to identify cell subpopulations from complex cell mixtures (Bakken *et al*, 2017). For instance, recent advances in single-cell RNA-sequencing (scRNA-seq) enable simultaneous measurement of expression levels of hundreds to thousands of genes across hundreds to thousands of individual cells. The resulting gene expression matrices of genes by cells are used (see below) to identify cell subpopulations with characteristic gene expression profiles and other biological properties (i.e. cell types).

A typical computational pipeline to process scRNA-seq data involves the following steps: *i*) quality control of sequencing reads, *ii*) mapping reads against a reference transcriptome, *iii*) normalization of mapped reads to correct batch effects and remove contaminants, *iv*) data dimensionality reduction with Principal Component Analysis or alternative approaches, *v*) clustering of cells using principal component values, *vi*) detection of genes differentially expressed between clusters, *vii*) visualization of cell clusters in t-SNE or alternative plots, and *viii*) assignment of cell type labels to cell clusters. A number of computational tools, including Cell Ranger (Zheng *et al*, 2017a) and Seurat (Butler *et al*, 2018), allow automation of steps *i* to *vii* (Innes & Bader, 2018; Freytag *et al*, 2018; Duò *et al*, 2018). However, assignment of cell type labels to cell clusters is still conducted manually by most researchers. The typical procedure involves manual inspection of the genes expressed in a cluster, combined with a detailed literature search to identify if any of those genes are known gene expression markers for cell types of interest. This manual approach has several caveats, including limited documentation and low reproducibility of cell type gene marker selection, use of uncontrolled and non-ontological vocabularies for cell type labels, and it can be time-consuming. For these reasons computational tools that allow researchers to systematically, reproducibly and quickly assign cell type labels to cell clusters derived from scRNA-seq experiments are needed.

In this study we used three scRNA-seq datasets from liver cells (MacParland *et al*, 2018), peripheral blood mononuclear cells (PBMCs) (Zheng *et al*, 2017a) and retinal neurons (Shekhar *et al*, 2016b) (Table 1) to compare four methods that can be used for assigning cell type labels to cell clusters: CIBERSORT (Newman *et al*, 2015b), GSEA (Subramanian *et al*, 2005), GSVA (Hänzelmann *et al*, 2013) and ORA (Fisher, 1935; Goeman & Bühlmann, 2007) (Table 2). We chose these four methods to represent different categories of methods, ranging from first-generation enrichment analysis (ORA) to second-generation approaches (GSEA and GSVA) and machine learning tools (CIBERSORT). Although ORA and GSEA were not originally developed to process RNA-seq data, they have been extensively used in transcriptomic studies for gene set enrichment analyses. GSVA was developed to analyse microarray and bulk RNA-seq data, and CIBERSORT was developed to estimate abundances of cell types in mixed cell populations from bulk RNA-seq data. We adapted all four methods to assign cell type labels to cell clusters from scRNA-seq data based on known sets of cell type marker genes. We evaluated these methods using two types of inputs: a matrix with the average expression of each gene *x* from all the cells in each cell cluster *y* (*Ě*_*xy*_) from scRNA-seq measurements, which we assume corresponds to the profile of a cell type or state, and known cell type gene expression signatures, represented as gene sets or continuous gene expression profiles (Figures 1A to 1C).

**Table 1.** scRNA-seq datasets used in this study. Details of the three datasets used in this study

| Dataset Name | Description of scRNA-seq dataset | Number of genes in $\tilde{E}_{xy}$ | Number of cells | Number of cell clusters | Number of cell type signatures | Reference |
| --- | --- | --- | --- | --- | --- | --- |
| Liver | 10X Chromium sample from liver cells from five human donors | 20,007 | 8,444 | 20 | 10 | (MacParland *et al*, 2018) |
| PBMCs | 10X Chromium sample from peripheral blood mononuclear cells from a human donor | 17,786 | 68,579 | 12 | 22 | (Zheng *et al*, 2017a) |
| Retinal neurons | Drop-seq sample from retinal bipolar neurons from healthy mice | 13,166 | 27,499 | 18 | 15 | (Shekhar *et al*, 2016b) |

**Table 2.** Cell cluster labeling methods compared in this study. Details of the four methods compared in this study, and their computing times to classify cell clusters of indicated datasets. (b) refers to CIBERSORT ‘binary’ analysis mode, (c) refers to CIBERSORT ‘continuous’ analysis mode.

| Acronym | Version | Name | Language | Computing time (s)* |  |  | Reference |
| --- | --- | --- | --- | --- | --- | --- | --- |
|  |  |  |  | liver | PBMCs | retinal neurons |  |
| CIBERSORT | 1.01 | <u>Cell type Identification by Estimating Relative Subsets of RNA Transcripts</u> | R and Java | (b) = 44 | (b) = 169<br>(c) = 700 | (b) = 36 | (Newman *et al*, 2015b) |
| GSEA | 3.0 | <u>Gene Set Enrichment Analysis</u> | Java | 93 | 78 | 98 | (Subramanian *et al*, 2005) |
| GSVA | 1.30 | <u>Gene Set Variation Analysis</u> | R | 1.2 | 0.9 | 0.73 | (Hänzelmann *et al*, 2013) |
| ORA | R(3.5.1) | <u>Over-representation Analysis</u> | R | 4 | 3 | 4 | (Fisher, 1935; Goeman & Bühlmann, 2007) |

**Figure 1.**
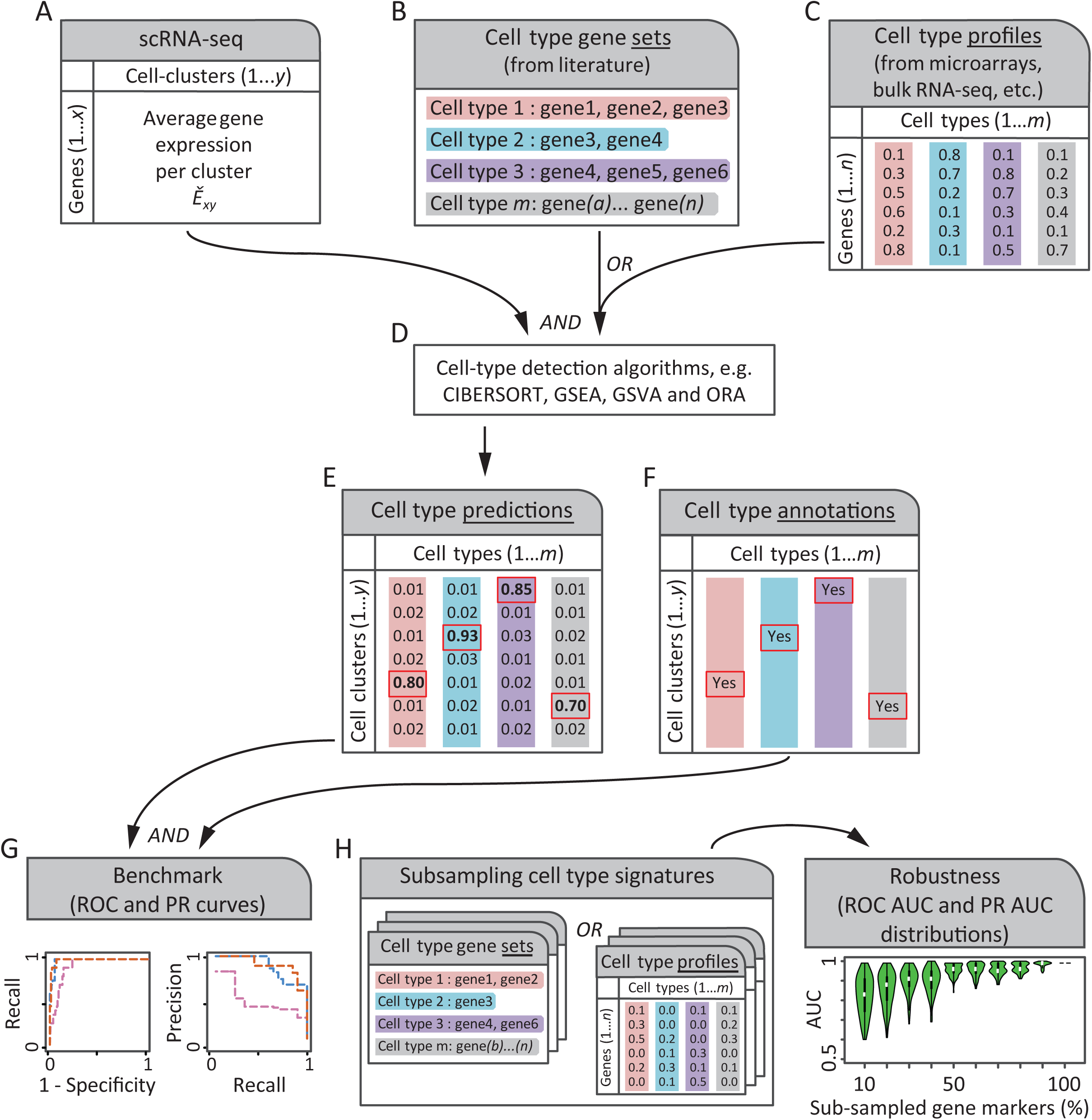
Schematic of a process to benchmark automated cell type detection methods. Two inputs are needed by automated cell type detection methods (panels A to C). **A.** a matrix with the average expression of each gene *x* for each cell cluster *y* (*Ě*_*xy*_). **B.** and **C.** cell type gene marker signatures can be provided as either gene sets (lists of gene identifiers, panel B) or numeric gene expression profiles (panel C). Gene sets can be manually compiled from literature and are used for methods like GSEA, GSVA or ORA. Whereas gene-expression profiles are measurements from microarrays, bulk-or scRNA-seq experiments, and are used by methods like CIBERSORT. **D.** and **E.** Automated cell type detection methods produce a matrix of cell type likelihoods for each cell cluster. **F.** Some authors of scRNA-seq studies assign cell type labels manually to cell clusters using empirical expertise or orthogonal experiments such as fluorescence activated cell sorting. These assignments can be used as references to benchmark automated cell type detections. **G.** Top cell type predictions (red rectangles in panel E) are contrasted against annotation references (panel F) to assess the performance of cell type detection methods by Receiver Operating Characteristic (ROC) curve and Precision-Recall (PR) curve analyses. **H.** Robustness of cell type detection methods can be analysed by gradually subsampling gene markers from cell type gene expression signatures (panels B or C) and repeating procedures of panels D to G to obtain distributions of the area under the curve (AUC) for ROC (ROC AUC) and PR (PR AUC) curves, which are shown as violin plots. We hypothesized that some detection methods may be more robust than others to the proportion of gene markers subsampled from cell type gene expression signatures.

CIBERSORT uses gene expression profiles as training data for a machine learning algorithm to estimate abundances of known cell types in a mixed cell population and was originally developed to identify composition of known immune cell types in bulk RNA-seq sample measurements. In our evaluation, we used *Ě*_*xy*_ matrices instead of bulk RNA-seq data. GSEA uses a Kolmogorov–Smirnov (KS) like statistic to determine whether a gene set shows statistically significant, concordant differences between biological states. It was originally developed to analyse microarray gene expression data and has been applied to multiple genomic data types. GSVA transforms a gene by sample matrix to a gene set by sample matrix, and evaluates gene set enrichment for each sample. Like GSEA, GSVA uses a KS like statistic but GSVA bypasses explicitly modeling phenotypes within the enrichment scoring step. GSVA was originally developed to process microarray and bulk RNA-seq measurements. ORA uses the Fisher’s exact test to detect an overrepresentation of members of a gene set in a subsample of highly expressed genes, compared against both the total number of gene set members and the total number of genes measured in the sample.

Methods explicitly developed to assign cell type labels to cell clusters of scRNA-seq data have been reported (Crow *et al*, 2018; Alquicira-Hernandez *et al*, 2018; Alavi *et al*, 2018). However, to our knowledge they are in beta, or implemented as web-servers to process cell types for which we could not find reference cell type annotations (Figure 1F) that we would require to include in our evaluation. For this reason, we included only the four methods described above, and we provide execution and benchmark scripts that will be useful to extend our comparisons to other methods in the future.

## Methods

### Generation of cell cluster average gene expression matrices (*Ě*_*xy*_)

For the liver dataset (MacParland *et al*, 2018) we followed the authors’ reported procedure to obtain cell clusters, and obtained the *Ě*_*xy*_ matrix for each cluster using the function AverageExpression(use.raw = T) from Seurat v2 (Butler *et al*, 2018). For the PBMCs dataset (Zheng *et al*, 2017a), Fresh 68k PBMCs DonorA gene expression matrix files were obtained from 10X (Zheng *et al*, 2017b). Normalization, data dimensionality reduction, cell clustering and *Ě*_*xy*_ matrix calculations were conducted with Seurat with the following functions: FilterCells(low.thresholds = 200,-Inf, high.thresholds = 0.05,10000); FindClusters(reduction.type = "pca", dims.use = 1:10, resolution = 0.4); AverageExpression(use.raw = T). For the retinal neurons dataset (Shekhar *et al*, 2016b) the gene expression matrix and cell cluster assignments were obtained from (Shekhar *et al*, 2016a) and the *Ě*_*xy*_ matrix calculation was conducted with AverageExpression(use.raw = T) from Seurat.

### Generation of cell type gene expression signatures

A gene expression signature is defined simply as a set of genes characteristically and detectably expressed in a cell type. These are typically identified in small scale experiments that need to be manually identified in the literature, or by comparing the transcriptome of a given cell type against all other available cell type gene expression profiles, usually from the same experiment. The liver cell type gene set signatures were manually curated by us (author S.A.M.) and were originally used to manually annotate cell types in the liver dataset (MacParland *et al*, 2018). We provide these gene sets at (Diaz-Mejia, 2019). For the PBMC dataset, we used a blood cell type gene expression profile signature compiled by the CIBERSORT developers called LM22, containing 547 genes and 22 cell types (Newman *et al*, 2015a). Reference cell type assignments to the PBMCs by fluorescence-activated cell sorting (FACS) were obtained from (Zheng *et al*, 2017c). The PBMC cell clusters we obtained with Seurat were mapped using cell barcode identifiers against the FACS assignments, and cell type names were manually matched to the LM22 signature. For the retinal neuron dataset (Shekhar *et al*, 2016b), known cell type markers reported by the authors were used as cell type gene set signatures.

CIBERSORT requires as input a cell type gene expression signature in the form of gene expression profiles (i.e. a matrix of genes in rows and cell types in columns). For the PBMC dataset, we used two versions of the LM22 signature for CIBERSORT. First, we used the original LM22 signature (Newman *et al*, 2015b) with continuous valued gene expression measurements, that we called CIBERSORT ‘continuous’. Second, for each cell type of the LM22 signature, a value of ‘1’ was assigned to 5% of genes with highest expression values in their column or a value of ‘0’ otherwise, and we called this approach CIBERSORT ‘binary’. The same 5% of genes was used to create cell type gene set signatures as inputs for GSEA, GSVA and ORA. For the liver dataset, we transformed the cell type gene set signature into a binary matrix of genes in rows and cell types in columns for CIBERSORT ‘binary’ analysis mode. To do this, each gene included in each cell type gene set *m* was assigned a value of ‘1’ in the column corresponding to *m* in the matrix, whereas other genes absent in *m* but present in other cell type gene sets were assigned a value of ‘0’. Similarly, for the retinal neuron dataset the ‘previously known markers’ for bipolar cell types provided in Table S2 of (Shekhar *et al*, 2016b) were transformed into a binary matrix of genes by cell types for CIBERSORT ‘binary’ analysis.

### Generation of subsampled cell type gene expression signatures and AUC violin plots

Cell type gene set signatures (Figure 1B) were subsampled by randomly removing between 10 and ∼99% of genes from each signature in increments of 10%, keeping a minimum of one gene. Each subsampling of gene sets was transformed into a binary matrix of genes by cell types for CIBERSORT ‘binary’ as indicated above. Cell type gene expression profile signatures (Figure 1C) were subsampled in two stages: first we selected the top 5% highest expressed genes for each cell type, then we randomly replaced the gene expression value of 10 to 100% of those genes from each cell type, in increments of 10%, by the minimum value of the cell type column. This resulted in subsampled gene expression profile signatures with identical size to the original profile signatures, but with values of the top highly expressed genes randomly replaced by the minimum score of each cell type. For percentage values between 10 to 100%, 1,000 subsampling replicates were generated for each cell type gene expression signature, and each replicate was processed as indicated by Figures 1D to 1G. Violin plots were used to show the resulting ROC and PR AUC distributions.

### Transformation of tested methods’ enrichment metrics for ROC and PR analyses

The enrichment scores (ES) from CIBERSORT and GSVA were directly used as ranks for the benchmark comparisons against gold standard references, whereas the P-values from GSEA and ORA were first -log 10 transformed and the resulting values were used as ranks for the benchmark analyses. For ORA, the universe of genes used was the intersection of genes present in the cell type gene expression signature and the *Ě*_*xy*_ matrix of each dataset. All methods were implemented locally using Java, R and Perl (Table 2) using the following libraries and programs: for CIBERSORT we used CIBERSORT.jar and R(Rserve), for GSEA we used gsea-3.0.jar, for GSVA we used R(GSVA) and R(GSA), and for ORA we used R(fisher.test).

### Method computing time benchmark

We implemented wrapper scripts to execute each of the four methods tested, including a stopwatch to time the cell type prediction task. Other tasks, such as input and output preparation, were excluded from computing time values reported in Table 2. All computing time measurements were made using a 3.1-GHz Intel Core i5 CPU with 2 cores and 16GB RAM.

## Results

We benchmarked the performance and computing time of four cell type labeling methods: CIBERSORT, GSVA, GSEA and ORA (Table 2) using average gene expression profiles of scRNA-seq cell clusters and known cell type gene expression signatures. We used three scRNA-seq datasets: liver cells (MacParland *et al*, 2018), PBMCs (Zheng *et al*, 2017a) and retinal neurons (Shekhar *et al*, 2016b) (Table 1). Each method used two inputs: an *Ě*_*xy*_ matrix with the average gene expression for each cell cluster (Figure 1A) and a cell type gene expression signature, represented as either a gene set or a gene expression profile. Three of the four methods tested (GSVA, GSEA and ORA) used cell type gene set signatures (Figure 1B), whereas CIBERSORT used cell type gene expression profiles either with continuous or binarized values (Figure 1C). Each method produced a matrix of cell type predictions (Figure 1D and E) which was compared to manually annotated cell type references (Figure 1F) to conduct Receiver Operating Characteristic (ROC) and Precision-Recall (PR) curve analyses (Figure 1G). The robustness of each method was assessed by randomly subsampling 10% to 100% of the genes from the cell type gene expression signatures and repeating the cell type detection and ROC and PR curve analyses for each subsample (Figure 1H).

In general, we observed that all four methods showed high ROC AUC values for all three analysed scRNA-seq datasets. An average ROC AUC = 0.97 was found for the liver dataset (Figure 2A), average ROC AUC = 0.92 for the PBMC dataset (Figure 2B) and average ROC AUC = 0.94 for the retinal neuron dataset (Figure 2C). Since CIBERSORT takes as input a cell type gene expression signature in the form of gene expression profiles (Figure 1C), and the only available signatures for the liver and retinal neuron datasets were in the form of gene sets, we transformed the gene sets into binary matrices and used them as inputs for CIBERSORT (Methods). Interestingly, the binary matrix approach, which we called CIBERSORT ‘binary’, produced the highest performance among all tested methods for the liver (ROC AUC = 1, Figure 2A) and retinal neurons datasets (ROC AUC = 0.95, Figure 2C). The CIBERSORT ‘binary’ approach performance was almost identical to that of the original LM22 cell type gene expression signature with continuous values, which we called CIBERSORT ‘continuous’, for the PBMC dataset (ROC AUC = 0.91 and 0.92, Figure 2B). GSVA was the top performer using the PBMC dataset (ROC AUC = 0.95, Figure 2B), closely followed by GSEA (ROC AUC = 0.94) and the two versions of CIBERSORT (‘binary’ ROC AUC = 0.92 and ‘continuous’ ROC AUC = 0.91), while ORA’s performance was slightly lower (ROC AUC = 0.86) (Figure 2B).

**Figure 2.**
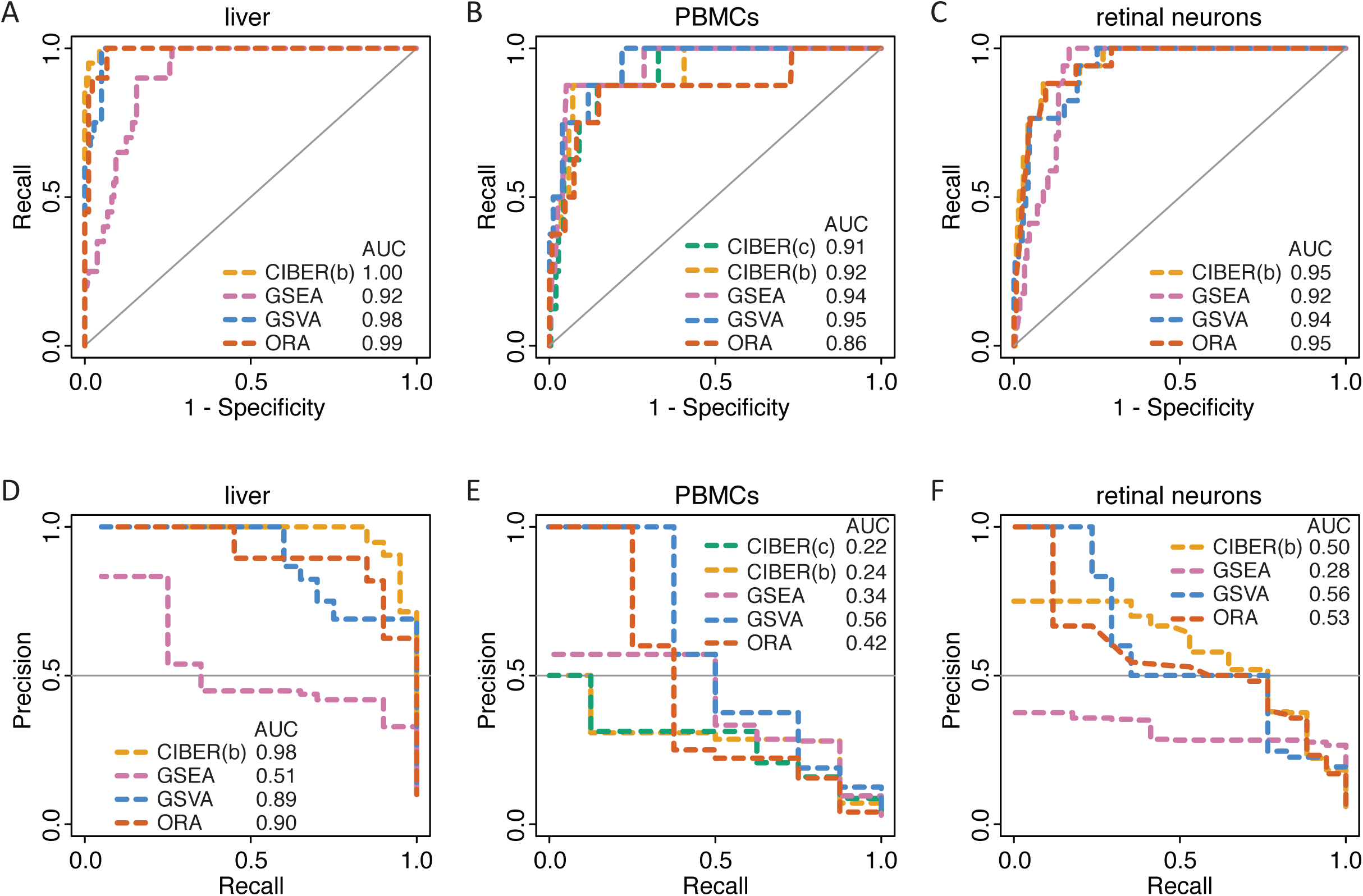
Performance analysis of automated cell type detection methods using scRNA-seq data. ROC and PR curve analyses of four automated cell type detection methods (CIBERSORT, GSEA, GSVA and ORA) (Table 2) using three scRNA-seq datasets (Table 1). ROC curve analyses for datasets from: **A.** human liver cells, **B.** human PBMCs, and **C.** mouse retinal neurons. PR curve analyses for the same datasets: **D.** human liver cells, **E.** human PBMCs, and **F.** mouse retinal neurons. The ROC AUC and PR AUC are shown for each method using each dataset. For the PBMCs dataset, two analyses were conducted with CIBERSORT, one using the original LM22 cell type gene expression signature with continuous gene expression values, that we called CIBERSORT ‘continuous’ (CIBER(c)), and another where the LM22 profiles were thresholded and binarized, that we called CIBERSORT ‘binary’ (CIBER(b), see Methods). The same thresholded signature was used to create cell type gene sets for GSEA, GSVA and ORA (Methods). For the liver and retinal neuron datasets only gene set signatures were available and they were transformed into binary matrices for CIBERSORT ‘binary’ (CIBER(b)).

The analysis of ROC AUC robustness showed that, in general, all methods’ performance decayed as a function of removing genes from cell type gene expression signatures. However, GSVA tolerated removal of up to 90% of the genes from the PBMC signature to maintain ROC AUC’s >= 0.8. ORA tolerated removal of up to 60% of genes at the same ROC AUC cutoff (Figure 3B), whereas GSEA and the two versions of CIBERSORT gave ROC AUC’s < 0.8 when >= 30% of the genes were removed from the PBMC cell type signatures. For the liver dataset, GSVA and GSEA tolerated removal of up to 60% of genes from the liver signature to maintain ROC AUC’s >= 0.8, whereas CIBERSORT ‘binary’ and ORA tolerated removal of up to 50% of the genes at the same ROC AUC cutoff (Figure 3A). For the retinal neuron dataset, GSVA and ORA tolerated removal of up to 50% of the genes from the signature to maintain ROC AUC’s >= 0.8, whereas GSEA and CIBERSORT ‘binary’ tolerated removal of 30% and 20%, respectively, for the same ROC AUC cutoff (Figure 3C).

**Figure 3.**
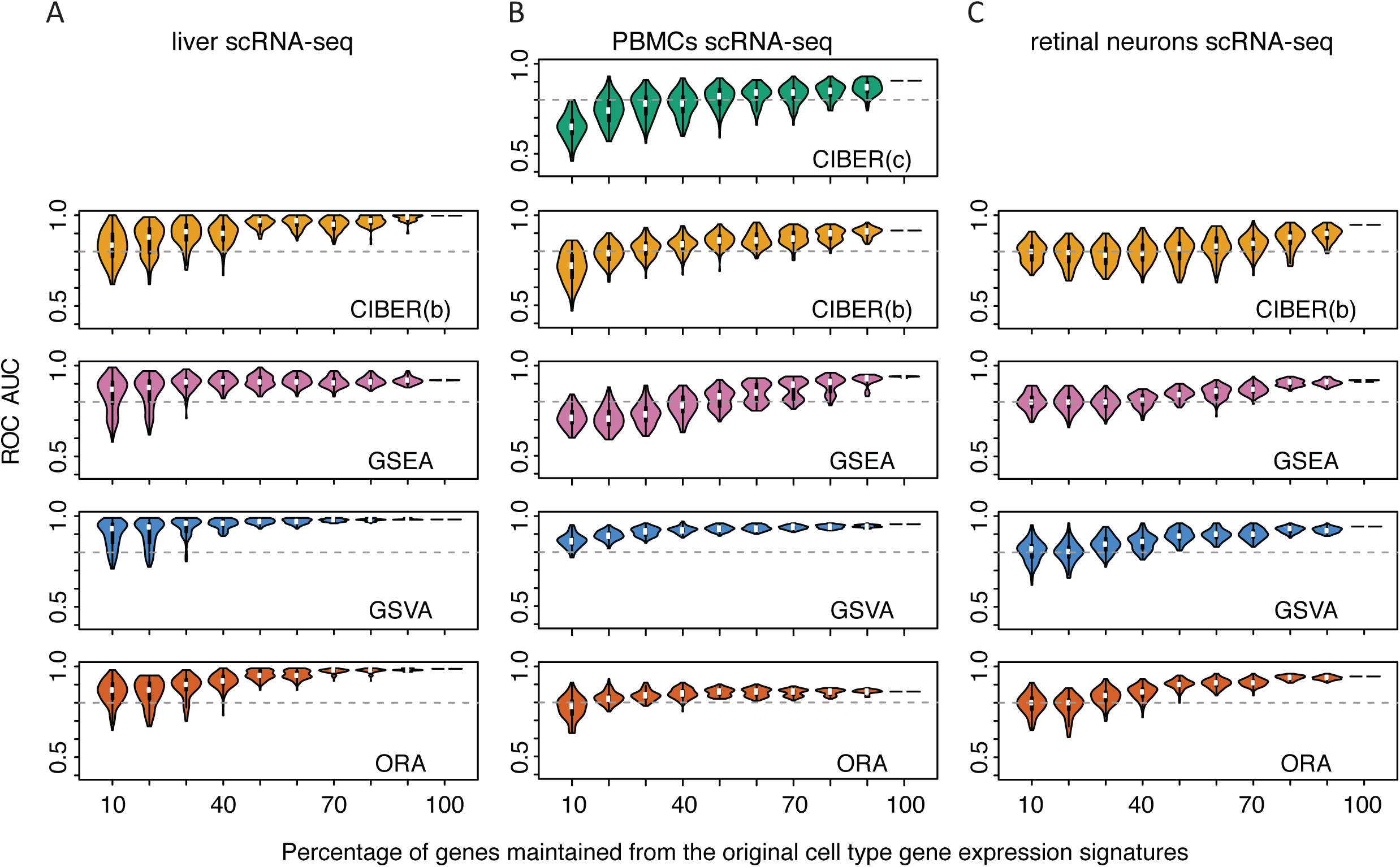
ROC AUC robustness analysis of automated cell type detection methods. The cell type gene expression signatures used for ROC curve analyses in Figure 2 were randomly subsampled 1,000 times, keeping 10 to 100% of genes from the original signatures. Automated cell type detection was repeated for each subsample and violin plots representing the distribution of resulting ROC AUC’s are shown for datasets from: **A.** human liver cells, **B.** human PBMCs, and **C.** mouse retinal neurons. For the PBMC dataset, two analyses were conducted with either the original LM22 cell type gene expression signature with continuous gene expression values (CIBER(c)) or with a thresholded and binarized version (CIBER(b)). For the liver and retinal neuron datasets only binary matrices for CIBER(b) were used.

When benchmarking the four methods compared in this study, we classified each cell cluster positively into a single cell type and negatively into the remaining cell types of their corresponding dataset signature. This produced a skewed distribution with few positive predictions and several negative predictions. To ameliorate this imbalance, we used PR curve analyses in addition to ROC curve analyses. In general, the PR AUC’s were smaller than the ROC AUC’s (Figure 2, top vs. bottom panels). Some methods clearly separated from the rest using PR curve analyses. For instance, GSEA showed the lowest PR AUC values for both the liver and retinal neurons datasets (PR AUC’s = 0.51 and 0.28), compared with CIBERSORT (PR AUCs = 0.98 and 0.5), ORA (PR AUC’s = 0.90 and 0.53), and GSVA (PR AUC = 0.89 and 0.56) (Figures 1D and 1F). GSEA also displayed the lowest AUC in the ROC curve analyses for the liver and retinal neurons datasets, and the performance differences between GSEA and the other methods were more pronounced using PR curve analyses. In contrast, the two versions of CIBERSORT for the PBMC dataset ranked very close to the other three methods using ROC curve analyses (all ROC AUC’s were > 0.9, Figure 2B), but they were relatively low using PR curve analyses (CIBERSORT ‘continuous’ PR AUC = 0.22 and CIBERSORT ‘binary’ PR AUC = 0.24), compared with GSVA (PR AUC = 0.56), ORA (PR AUC = 0.42) and GSEA (PR AUC = 0.34) (Figure 2E).

The PR AUC robustness analysis showed that all methods’ performance decayed as a function of removing genes from cell type gene expression signatures. Interestingly, using the liver dataset all four methods showed higher PR AUC’s than for the PBMC and retinal neuron datasets (Figure 4A vs. 4B and 4C). In particular, GSVA and ORA tolerated removal of up to 60% of genes from the liver dataset signatures to maintain PR AUC’s >= 0.5. CIBERSORT ‘binary’ tolerated removal of 50% of genes for the same PR AUC cutoff (Figure 4A), whereas GSEA PR AUC’s were < 0.5 using either the full PBMC cell type signature or any subsampling of it. For the retinal neuron dataset, CIBERSORT ‘binary’, GSVA and ORA tolerated removal of up to 20% of the genes from the signature to maintain average PR AUC >= 0.5, whereas for GSEA the average was < 0.5 at any fraction of genes in the signature. For the PBMC dataset, GSVA was the only method showing PR AUC > 0.5 with the full signature (Figure 2E) and it tolerated removal of up to 20% of genes from the signature to maintain average PR AUC > 0.5 (Figure 4B).

**Figure 4.**
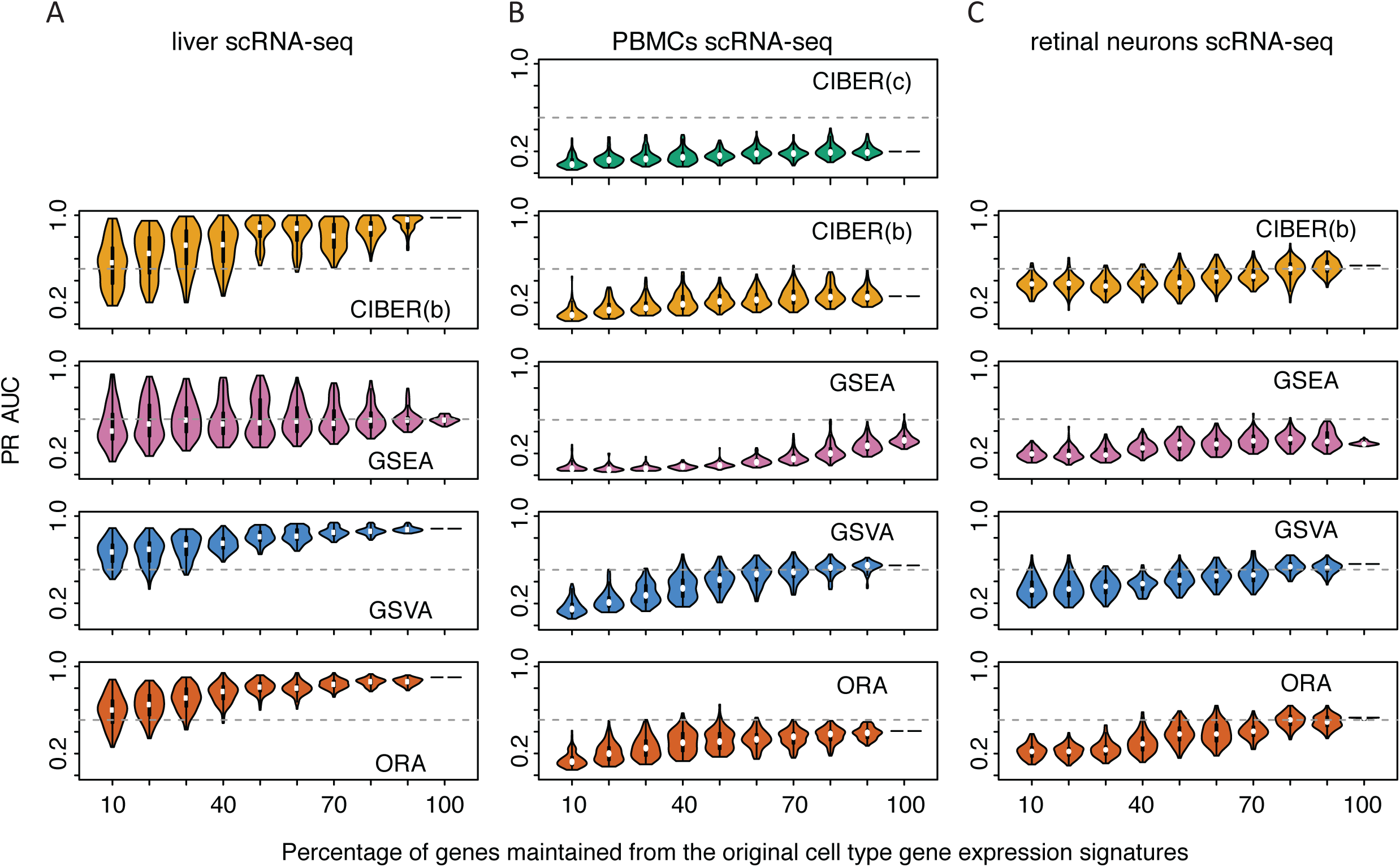
PR AUC robustness analysis of automated cell type detection methods. The same procedure described in Figure 3 for ROC AUC’s was used here for PR AUC’s. Please see Figure 3 legend for details.

As shown in Table 2, the computing times of method implementations varied from 0.73s for GSVA processing the retinal neurons dataset, up to 700 s for CIBERSORT ‘continuous’ processing the PBMC dataset. For all three datasets, GSVA was the fastest method to process cell type classification tasks. ORA ranked second with computing times between 3 and 5 times longer than GSVA. GSEA showed computing times between 77 and 134 times longer than GSVA, and CIBERSORT showed computing times between 37 and 777 times longer than GSVA. The size of the cell type gene expression signatures used for CIBERSORT influenced the speed of the classification task. For CIBERSORT ‘continuous’ we used the original LM22 signature, which contained 547 genes for the PBMC dataset, whereas the thresholded binary matrix used for CIBERSORT ‘binary’ had 248 genes, and it took 169s, or 24% ot the time that took CIBERSORT ‘continuous’ for the same task. For comparison, we created a second ‘continuous’ signature by restricting the original LM22 signature to the 248 genes present in the thresholded binary matrix. This ‘reduced continuous’ signature approach showed a performance (ROC AUC = 0.92, PR AUC = 0.32) which was similar to the full CIBERSORT ‘continuous’ (ROC AUC = 0.92, PR AUC = 0.24) and ‘binary’ modes (ROC AUC = 0.91, PR AUC = 0.22), and the computing time was reduced substantially to 189s, or 27% of the time that took CIBERSORT ‘continuous’ for the same task.

## Discussion

The size and volume of scRNA-seq datasets are continually increasing, and several methods are available to normalize scRNA-seq measurements and cluster cells. In contrast, cell type labeling of cell clusters is still conducted manually by most researchers. This is in part due to a scarcity of reference cell type gene expression signatures and also because most methods to address this challenge are only available via web servers with limited number of cell types (Crow *et al*, 2018; Alquicira-Hernandez *et al*, 2018; Alavi *et al*, 2018), making it difficult for users to adapt them for their needs. In this study we used three scRNA-seq datasets to benchmark four methods that can address these challenges. Although three of the four tested methods (GSEA, GSVA and ORA) were not explicitly developed to identify cell types, their extensive use in gene set enrichment tasks and their wide portability motivated us to test them as cell type classifiers. CIBERSORT is implemented both as a webserver and a local distribution can be licensed by developers, allowing use to benchmark it with relatively low programmatic effort.

In general, our results show that for the three scRNA-seq datasets tested (liver, PBMCs and retinal neurons) all four tested methods achieved good performance by ROC curve analyses. However, ROC curves tend to overestimate methods’ performance when the ratio of positive to negative predictions is highly skewed. For this reason, we decided to also conduct PR curve analyses. GSVA was consistently one of the top performers by both ROC and PR curve analyses for the three datasets, and its performance was more robust in analyses where we subsampled genes from cell type gene expression signatures. This is particularly important at this stage of the scRNA-seq field, as only limited information on cell type gene expression signatures is available. Interestingly, despite its relative simplicity, ORA showed a performance comparable to GSVA. CIBERSORT’s performance was good, particularly for the liver dataset by both ROC and PR analyses, albeit lower than that of GSVA or ORA in the PBMC dataset, and it was comparable using the retinal neuron dataset. CIBERSORT’s computing times were orders of magnitude higher those of GSVA and ORA. Our results showed that CIBERSORT ‘binary’ performed as well as CIBERSORT ‘continuous’ by both ROC and PR curve analyses and used only one quarter of the computing time. In the present implementation, GSEA performed worse than the other three methods, particularly in the PR curve analyses.

The size of current publicly available scRNA-seq datasets is currently typically on the order of thousands of cells clustered into dozens of cell clusters. In our tests, each of the four tested methods completed the cell type prediction tasks in seconds or minutes. However, bigger datasets from the Human Cell Atlas (Rozenblatt-Rosen *et al*, 2017) and other sources are expected to have millions of cells (e.g. https://support.10xgenomics.com/single-cell-gene-expression/datasets/1.3.0/1M_neurons) grouped into thousands of clusters, for which the fastest method implementations will be preferred. In this sense, we found that GSVA is the best option since its computing time for the tested datasets was fastest (one to two orders of magnitude faster than GSEA and CIBERSORT). ORA also offers a good option for cell cluster labeling as its ROC and PR curve benchmarks were comparable to GSVA and its computing times were only 3 to 5 times longer than those of GSVA. One extra requirement for ORA compared with the other three methods is that the *Ě*_*xy*_ matrix profiles need to be thresholded. In this study we used an arbitrary cutoff, based on the overall distribution of gene expression values, but future analyses could evaluate iterative thresholding.

One of the limitations of this study is that we included only three scRNA-seq datasets (liver, PBMCs and retinal neurons). This was due to the lack of reference cell type annotations needed for the ROC and PR curve analyses. As more scRNA-seq datasets become available and authors provide gold standard annotations of their cell types, those annotations could be used as references to benchmark methods with other scRNA-seq datasets. This is exemplified by the LM22 signature, which was constructed by (Newman *et al*, 2015b) from microarray gene expression measurements to predict cell types from bulk RNA-seq data, and we have shown here that LM22 could also be used to detect cell types from scRNA-seq data. Thus, in the future, we envision that methods to detect differentially expressed genes can be used as part of pipelines to produce cell type gene expression signatures. As with any classification task, researchers would need to control for circularity between training, test and validation cell-annotation data and also will need to evaluate generalizability.

One of the challenges that we faced while adapting the LM22 signature to detect cell types in the scRNA-seq cell clusters from (Zheng *et al*, 2017a) was that, even though both datasets correspond to PBMCs, the granularity of their cell type labels was different. For instance, the LM22 signature contains six T-cell types, including three CD4+ (naïve, memory resting, and memory activated), follicular helper, regulatory and gamma delta, whereas the (Zheng *et al*, 2017a) dataset contained labels for four T-cell related cell types: CD4+/CD25 T Regulatory, CD4+/CD45RO+ Memory, CD4+/CD45RA+/CD25-Naive T and CD4+ T Helper2. Thus, even though these two datasets both classify PBMCs, they cannot be easily related one-to-one. This could be addressed with an ontology analogous to the Gene Ontology (Ashburner *et al*, 2000) but dedicated to cell type annotations (Bard *et al*, 2005; Bakken *et al*, 2017). Fortunately, the Cell Ontology is being developed for this purpose (https://www.jcvi.org/cl-cell-ontology). This is particularly important as an increasing number of signatures are expected to arise from initiatives like the Human Cell Atlas (Rozenblatt-Rosen *et al*, 2017).

## Data and Software Availability

We provide our R and Perl scripts used to run and benchmark cell type labeling methods to help researchers to reproduce our analyses, or to test other scRNA-seq datasets or cell type labeling method implementations, at Github: https://github.com/jdime/scRNAseq_cell_cluster_labeling.

The three scRNA-seq datasets from liver cells liver cells (MacParland *et al*, 2018), PBMCs (Zheng *et al*, 2017a) and retinal neurons (Shekhar *et al*, 2016b), were deposited at Zenodo: “Evaluation of methods to assign cell type labels to cell clusters from single-cell RNA-sequencing data” at http://doi.org/10.5281/zenodo.2575050

## Author Contributions

JJDM: Conceptualization, Data Curation, Analysis, Software, Supervision, Writing Draft & Editing
EM: Data Curation, Writing Draft & Editing
ARP: Conceptualization
SAM: Data Curation, Writing Draft & Editing
TK: Writing Draft & Editing
TJP: Writing Draft & Editing
GDB: Conceptualization, Supervision, Writing Draft & Editing
JHM: Conceptualization, Supervision, Writing Draft & Editing

## Acknowledgements

We would like to thank to Jeff Liu and Brendan Innes from the Bader lab for advice processing the liver dataset and implementing GSVA; to Danielle Croucher and Laura Richards from the Pugh lab for feedback collecting benchmark datasets; and to Rene Quevedo from the Pugh lab for help implementing R scripts.

